# Implantable flexible multielectrode arrays for multi-site sensing of serotonin tonic levels

**DOI:** 10.1101/2023.01.17.524488

**Authors:** Elisa Castagnola, Elaine M. Robbins, Daniela Krahe, Bingchen Wu, May Yoon Pwint, Qun Cao, Xinyan Tracy Cui

## Abstract

Real-time multi-channel measurements of tonic serotonin (5-hydroxytryptamine, 5-HT) concentrations across different brain regions are of utmost importance to the understanding of 5-HT’s role in anxiety, depression, and impulse control disorders, which will improve the diagnosis and treatment of these neuropsychiatric illnesses. Chronic sampling of 5-HT is critical in tracking disease development as well as the time course of pharmacological treatments. Despite their value, *in vivo* chronic multi-site measurements of 5-HT have not been reported.

To fill this technological gap, we batch fabricated implantable glassy carbon (GC) microelectrode arrays (MEAs) on a flexible SU-8 substrate to provide an electrochemically stable and biocompatible device/tissue interface. Then, to achieve multi-site detection of tonic 5-HT concentrations, we incorporated the poly(3,4-ethylenedioxythiophene)/functionalized carbon nanotube (PEDOT/CNT) coating on the GC microelectrodes in combination with a new square wave voltammetry (SWV) approach, optimized for selective 5-HT measurement. *In vitro*, the PEDOT/CNT coated GC microelectrodes achieved high sensitivity towards 5-HT, good fouling resistance in the presence of 5-HT, and excellent selectivity towards the most common neurochemical interferents. *In vivo*, our PEDOT/CNT-coated GC MEAs were able to successfully detect basal 5-HT concentrations at different locations of the CA2 hippocampal region of mice in both anesthetized and awake head-fixed conditions. Furthermore, the implanted PEDOT/CNT-coated MEA achieved stable detection of tonic 5-HT concentrations for one week. Finally, histology data in the hippocampus shows reduced tissue damage and inflammatory responses compared to stiff silicon probes. To the best of our knowledge, this PEDOT/CNT-coated GC MEA is the first implantable flexible multisite sensor capable of chronic *in vivo* multi-site sensing of tonic 5-HT. This implantable MEA can be custom-designed according to specific brain region of interests and research questions, with the potential to combine electrophysiology recording and multiple analyte sensing to maximize our understanding of neurochemistry.

**Highlights:**

- PEDOT/CNT-coated GC microelectrodes enabled sensitive and selective tonic detection of serotonin (5-HT) using a new square wave voltammetry (SWV) approach
- PEDOT/CNT-coated GC MEAs achieved multi-site *in vivo* 5-HT tonic detection for one week.
- Flexible MEAs lead to reduced tissue damage and inflammation compared to stiff silicon probes.

## 1. Introduction

Serotonin (5-hydroxytryptamine, 5-HT) plays a central role in a variety of brain functions, including cognition (Amin et al. 2005; Bacqué-Cazenave et al. 2020; Buhot 1997), emotion (Lesch 2005; Meneses and Liy-Salmeron 2012; Merens et al. 2007), and social behaviors (Crockett et al. 2008; Kiser et al. 2012; Lesch 2007). Deficiency of the serotoninergic system has been implicated in anxiety (Albert et al. 2014; Deakin 1998; Hammack et al. 2009), depression (Albert et al. 2014; Deakin 1998; Fakhoury 2016; Hammack et al. 2009; Neumeister et al. 2004; Yagishita 2020), and impulse control disorders (Krakowski 2003; Linnoila et al. 1993). 5-HT release occurs over multiple timescales (Cohen et al. 2015; Cools et al. 2008; Yagishita 2020), including phasic release (milliseconds to seconds, caused by neuronal firing in response to stimuli) (Cools et al. 2008) and tonic release (slow-changing basal levels on the seconds to minute timescale) (Cools et al. 2008). The alterations of both of these concentration paradigms have been found to result in abnormal neurological function (Cools et al. 2008; Yagishita 2020) and behavior (Cools et al. 2008; Gonzalez et al. 2004; Yagishita 2020). Measuring 5-HT signaling at different time scales will allow us to determine the different roles phasic and tonic 5-HT play, investigate the interplay between them, and shed light on the mechanisms behind 5-HT functions. Unfortunately, no sensor has been reported on chronic measurement of tonic or phasic 5-HT from the brain.

Traditionally, tonic 5-HT levels have been measured using microdialysis (Bungay et al. 2003; Guida et al. 2020; Zestos and Kennedy 2017). However, microdialysis suffers from poor temporal resolution (minutes (Yang et al. 1998)), and substantial tissue damage that greatly diminishes extraction efficiency over time (Beyene et al. 2019; Di Chiara et al. 1996; Jaquins-Gerstl and Michael 2015). N-shaped fast scan cyclic voltammetry (N-FSCV) detection of phasic 5-HT is well-established in acute *in vivo* studies, but chronic sensing over days has not been achieved (Carl J. Meunier 2019; Hashemi et al. 2009; Saylor et al. 2019). While FSCV is effective at measuring phasic 5-HT release at fast temporal resolutions (hundreds of milliseconds (Hashemi et al. 2009; Jacobs et al. 2014; Keithley et al. 2011; Robinson et al. 2003; Saylor et al. 2019)), the necessity for background subtraction has prevented its use for tonic 5-HT measurement (Carl J. Meunier 2019; Mark DeWaele 2017). Modified voltammetry techniques based on FSCV, i.e. fast-scan controlled-adsorption voltammetry (FSCAV) (Abdalla et al. 2017; Saylor et al. 2019), or multiple square waveforms superimposed onto an N-shaped cyclic waveform, i.e. N-Shaped Multiple Cyclic Square Wave Voltammetry (N-MCSWV) (Shin et al. 2021), have been developed with the attempt to enable 5-HT tonic detection. These techniques rely on the adsorption properties of the serotonin onto the surface of the CFEs and require the application of dynamic background subtraction (Shin et al. 2021), not entirely successful in eliminating non-faradaic components due to the fast scan rate and/or the capacitive charging current changes among CSWs. Thus, further mathematical exponential decay equation analysis, or statistical modeling are required for the quantification of the 5-HT faradaic currents (Abdalla et al. 2017; Shin et al. 2021).

Furthermore, these experiments are typically performed at carbon fiber microelectrodes (CFEs), one site at a time (Abdalla et al. 2017; Hashemi et al. 2009; Saylor et al. 2019; Yoonbae Oh 2016), while 5-HT dynamics are complex and differ in different brain regions or different loci of the same region, requiring high resolution multisite measurements (Kim et al. 2005; Perez-Garcia and Meneses 2008). In the last few years, GC MEAs have been introduced as a multimodal platform for integrated multi-channel electrophysiological and electrochemical measurements for neural applications (Castagnola et al. 2022; Castagnola et al. 2021b; Castagnola et al. 2018; Devi et al. 2021; Nimbalkar et al. 2018; Vahidi et al. 2020; Vomero et al. 2017). GC microelectrodes enable the integration of FSCV electrochemical sensing (Castagnola et al. 2022; Castagnola et al. 2021b; Castagnola et al. 2018) to devices that are routinely used for multi-channel neurophysiological recordings and stimulation. However, although GC electrodes provides a superior electrochemical stability (Nimbalkar et al. 2018; Vomero et al. 2017) and improved coating adhesion (Castagnola et al. 2017; Vomero et al. 2017) than metal microelectrodes, its effective surface area is lower than conducting polymers and nanocarbon. Poly(3,4-ethylenedioxythiophene)/acid functionalized carbon nanotube (PEDOT/CNT) is a high surface area electrode coating that has been shown to dramatically reduce impedance, improve electrophysiological recording (Alba et al. 2015; Kozai et al. 2015) and stimulation (Kolarcik et al. 2014; Luo et al. 2011; Zheng et al. 2022), and enable sensitive detection of tonic dopamine (DA) using SWV (Castagnola et al. 2022). In this work, we focused on the development for direct tonic 5-HT measurement using an optimized SWV waveform in combination with PEDOT/CNT coating on GC MEA.

For *in vivo* chronic measurements, healthy device-tissue interface is fundamentally important for high fidelity and stable recording (Edell et al. 1992; Kozai et al. 2015) and sensing over time (Xu et al. 2019). Stiff implants cause significantly lower neuronal density and increased glia encapsulation around the implant site (Edell et al. 1992; Kozai et al. 2015; Szarowski et al. 2003), while flexible implantable MEAs based on thin-film polymer substrate, matching the soft nature of the brain, can reduce mechanical micro-traumas and inflammation (He et al. 2020; Luan et al. 2020; Seymour and Kipke 2006; Seymour et al. 2017; Vomero et al. 2022; Xie et al. 2012). Several studies suggested that reduced mechanical mismatch between the flexible probe and brain tissue contributes to improved longevity of electrophysiological recording (He et al. 2020; Luan et al. 2017; Vomero et al. 2022; Xie et al. 2012; Zhao et al. 2019). Using a pattern-transfer technique (Castagnola et al. 2022; Castagnola et al. 2021b; Vomero et al. 2017), in combination with the high-resolution mask-less photolithography process (Castagnola et al. 2022), we integrated GC microelectrodes on SU-8 flexible substrate, and examine the acute and chronic 5-HT sensing capability of our flexible PEDOT/CNT-coated GC MEAs microelectrodes, as well as the host tissue response in comparison to stiff counterpart.

## 2. Material and Methods

### 2.1 Glassy Carbon Microelectrode Array Fabrication

Similar to what we previously reported (Castagnola et al. 2022), a 4-inch Si wafer with a 1 µm thick SiO2 layer (University Wafer Inc.) was first cleaned with acetone, isopropanol, and deionized (DI) water sequentially. The wafer was then dried with a N2 spray gun, heated on a hot plate at 150°C for 5 mins, and treated by O2 plasma using a reactive ion etcher (RIE, Trion Phantom III LT) for two minutes at 200 mTorr pressure and 150 W power. The cleaned wafer was spin-coated with SU-8 100 (MicroChemicals) at 6000 rpm for one minute and soft baked at 65°C for 5 mins and 95°C for 15 mins. Then the wafer was exposed to 365 nm ultraviolet (UV) light using a direct writing mask-less aligner (MLA, MLA100 Heidelberg Instruments) with a dose of 6000 mJ/cm2. After UV exposure, the wafer was first post baked at 65°C for 3 mins and 95°C for 8 mins, then developed using SU-8 developer (MicroChemicals) for four minutes and cleaned by isopropanol and DI water. The patterned SU-8 was subsequently hard baked at 200°C, 180°C, and 150°C for five minutes each, and allowed to cool down below 95°C. Pyrolysis of the negative SU-8 resist was performed at 1000 °C in an inert N2 environment for 60 minutes at 0.8 Torr.

After the pyrolysis step, the wafer was cleaned with acetone, isopropanol, and DI water sequentially, and treated with O2 plasma with RIE for 90 seconds at pressure of 200 mTorr and 150 W power. The cleaned wafer was then spin-coated with SU-8 2005 (MicroChemicals) at 4000 rpm for 1 min, and subsequently soft baked at 65°C for 3 mins and 95°C for 5 mins. Then the first SU-8 layer was patterned, using the MLA with a dose of 300 mJ/cm2, to define the bottom insulation layer and to open a connection between GC electrodes and the metal traces (next step). After a post bake at 65°C for 3 mins and 95°C for 5 mins, the wafer was developed using SU-8 developer. Finally, the patterned wafer was cleaned by isopropanol and DI water and hard baked at 200°C, 180°C, and 150°C for five minutes each, and allowed to cool down below 95°C.

After cleaning, the wafer was spin coated with AZ P4620 photoresist (MicroChemicals) at 5300 rpm for 1min and baked at 105°C for 5 mins. After soft baking, the wafer was exposed using MLA with a dose of 700 mJ/cm2, then developed using AZ400k 1:4 developer (MicroChemicals), rinsed with water, and dried by N2 gas flow. A 10nm Ti adhesion layer and 100 nm Au layer were evaporated on the wafer using an Electron Beam Evaporator (Plassys MEB550S) and then the metal was lifted-off in acetone to define the metal traces and connection pads. A top insulation layer of SU-8 2005 was then spin-coated at 4000 rpm for one minute, soft baked at 65°C for 3 mins and 95°C for 5 mins and photo-patterned, using the MLA with a dose of 300 mJ/cm2 to expose the connection pads and to define the top insulation layer. After post baking and development with the SU-8 developer, the wafer was cleaned with isopropanol and DI water, and hard baked at 200°C, 180°C, and 150°C for 5 mins each and allowed to cool down below 95°C. The MEAs were lifted-off from the wafer using buffered oxide etchant (1:7) in an acid hood for 4-6 hours. An anchor hole was also patterned at the shank tip to facilitate the insertion of a 50 µm tungsten shuttle, to enable the handling and penetration of the flexible device into the brain. **Figure SI 1 of Supplementary Information** shows the schematic of the GC-MEA fabrication.

### 2.2 Raman and SEM characterizations

Raman spectroscopy measurements were performed using Renishaw inVia Micro-Raman Microscope system (Renishaw, Hoffman Estates, IL) A 1800 lines/mm diffraction grating was used. A 633 nm laser was focused to a spot size on the sample through a 100x objective. Laser intensity was 100% and the scan range was 100 cm^-1^ to 3200 cm^-1^. Before all the measurements, the system was calibrated with Si (100) Reference Sample. All the data shown in Figure 1B were five-point averages collected from the same sample. Scanning electron microscope (SEM) is performed using FEI Scios Dual Beam System (Thermo Fisher Scientific, Waltham, MA) with following parameters: 5kV, 0.1nA beam. 7mm working distance.

**Figure 1.**
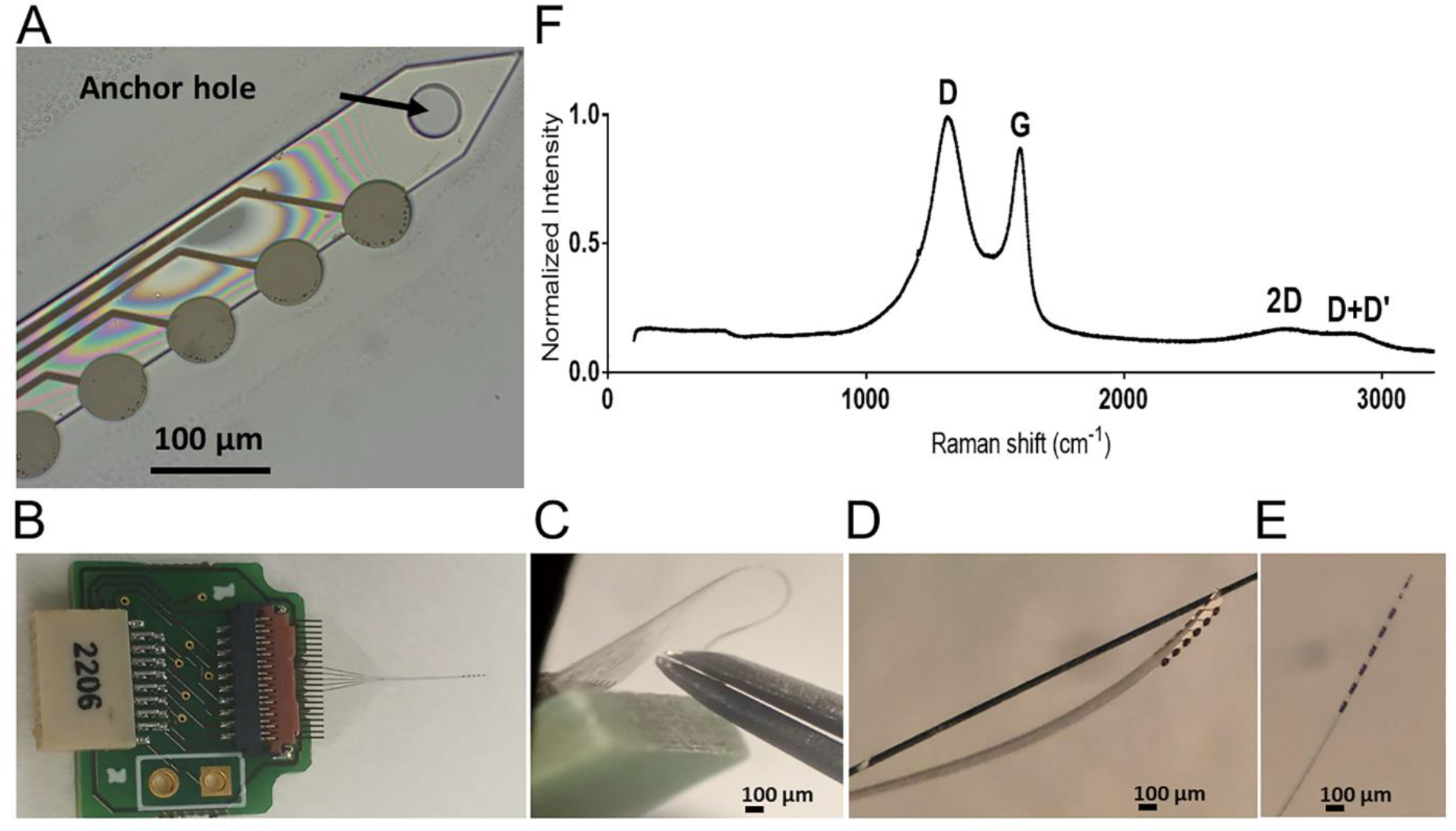
Flexible Glassy Carbon microelectrode arrays (GC-MEAs). **A)** MEA microfabricated with metal interconnection and GC microelectrodes, after release from the wafer. The GC-MEAs has been fabricated with an anchor hole at the shank tip to facilitate the insertion of a sharpened 50 µm tungsten wire, that enables the handling and penetration of the probes into the brain. **B, C, D, E)** Pictures of a flexible MEAs connected to the PCB using a zero-insertion force (ZIF) connector **(B, C)**, and assembled with the 50 µm tungsten wire **(D). E)** Optical picture the lateral view of the GC MEA. **F)** Raman spectra of the glassy carbon microelectrodes.

### 2.3 PEDOT/CNT coating and electrochemical characterization

Multi-walled CNTs (length of 10–30 μm and diameter of 20–30 nm, Cheap Tubes Inc.) were functionalized using our previously established methods (Kozai et al. 2016; Luo et al. 2011). PEDOT/CNT coatings were electropolymerized on the GC electrodes using a previously reported procedure (Kozai et al. 2016; Luo et al. 2011; Taylor et al. 2019). Briefly, electropolymerization was carried out in an aqueous solution of 0.02 M 3,4-ethylenedioxythiophene (EDOT; Sigma) containing 1 mg mL^−1^ of functionalized CNT with a constant potential of 0.9 V until a 120 mC/cm^2^ charge density cut-off was reached.

Electrochemical impedance spectroscopy (EIS) measurements were used to investigate the electrode/electrolyte interface before and after the PEDOT/CNT coating and to quantify their impedance in the 1 Hz -100 kHz range (Rose and Robblee 1990) *in vitro*, and to verify the functionality of the MEAs immediately after implantation, *in vivo*, as previously reported (Castagnola et al. 2020a; Taylor et al. 2019). During the EIS measurements, a sine wave (10 mV RMS amplitude) was superimposed onto the open circuit potential while varying the frequency from 1 to 10^5^ Hz.

Cyclic voltammetry (CV) was performed to quantify the capacitive charging of the GC microelectrodes before and after PEDOT/CNT coatings. During the CV tests, the working electrode potential was swept between 0.8 and −0.6 V vs. Ag/AgCl at a scan rate of 100 mV/s. The *in vitro* EIS and CV were performed in 1x PBS in a three-electrode electrochemical cell set-up with a platinum counter electrode and an Ag/AgCl wire reference electrode. *In vivo*, a bone screw was used as counter electrode and the Ag/AgCl wire reference electrode was placed in contact with the brain through a small pinhole craniotomy.

Electropolymerization, EIS and CV were carried out using a potentiostat/galvanostat (Autolab, Metrohm, USA).

### 2.4 SWV *in vitro* testing

Electrochemical detection of 5-HT was performed using an optimized SWV waveform. SWV experiments were carried out using a potentiostat/galvanostat (Autolab, Metrohm, USA) connected to a three-electrode electrochemical cell with a platinum counter electrode and an Ag/AgCl reference electrode. The SWV waveform was repeatedly applied from 0.15 V to 0.5 V with a 40 Hz step frequency, a 50 mV pulse amplitude and a 5 mV step height every 15 seconds. Potential was held at 0.15 V between scans. For each electrode used in vivo, preimplant calibration (pre-calibration) was performed. *In vitro* calibrations were performed using freshly prepared 5-HT solutions dissolved in artificial cerebrospinal fluid (aCSF) (142 mM NaCl, 1.2 mM CaCl2, 2.7 mM KCl, 1.0 mM MgCl2, 2.0 mM NaH2PO4, pH 7.4) in a 50 nM –1 μM concentration range. Electrode sensitivity was determined by the slope of the linear range of the calibration plot relating 5-HT peak current at 0.29 V to 5-HT concentration.

### 2.5 *In vivo* Procedures

The *in vivo* performance was determined through acute and chronic surgical experiments conducted in the CA2 region of the hippocampus of 8 male mice (C57BL/6J, 8–12 weeks, 22–35 g; Jackson Laboratory, Bar Harbor, ME). All animal care and procedures were performed under the approval of the University of Pittsburgh Institutional Animal Care and Use Committee.

All animals were induced with 1.5–2% isoflurane mixed with oxygen flow at 1 L min−1, then maintained at 1.25–1.5%. The body temperature was maintained at 37 °C with a thermostatically controlled heating pad (Harvard Apparatus, Holliston, MA, USA), and lacrigel (Dechra Puralube Vet Ointment) was placed on eyes to avoid dryness.

After the animal head was fixed in a stereotaxic frame (Narishige International USA, Inc), the skin and connective tissue on the surface of the skull was removed. A small pinhole craniotomy was made over the CA2 region of the hippocampus (AP: -2.91, ML: +3.35, DV: -2.5 to -3.0 relative to bregma)(Paxinos and Franklin 2019) with a high-speed dental drill (0.007 drill bit, Fine Science Tools, Inc., Foster City, CA) and bone fragments were carefully removed with forceps and saline. Saline was applied continuously onto the skull to dissipate heat from the high-speed drill.

The MEAs were lowered 3 mm below the cortical surface, into the CA2 using a hand-driven micromanipulator such that the 6 electrodes should be positioned in the CA2. Two additional small pinhole craniotomies were performed for the introduction of the Ag/AgCl reference electrode contralaterally to the MEA and a bone screw counter electrode caudally to the reference. EIS was measured immediately after the MEA implantation. Then, the tonic 5-HT response was measured using the SWV waveform (detailed above) over a ten-minute period sequentially for each microelectrode of the MEA. To confirm the chemical specificity of our measurements, the validation of the 5-HT peak was performed using stereotaxic direct infusion of 1 μL bolus of 5 and 10 µM 5-HT solutions. Specifically, in acute experiments, a fused silica capillary (75 μm ID, 150 μm OD, Polymicro Technologies, Phoenix, AZ, United States) was glued to the back side of the MEAs and the direct infusion was performed using a gastight syringe (1000 Series Gastight, Hamilton, Reno, NV, United States) driven by a syringe pump (Harvard Apparatus Model 11 (55-1199), Holliston, MA, United States). In acute experiments, upon reaching the predetermined experimental endpoint, the MEAs were explanted, and the animals were humanely sacrificed using approved procedures.

For the SWV chronic and head fixed awake experiments, after the PEDOT/CNT coated GC MEA positioning, the craniotomy was filled with Kwik-Cast Sealant (World Precision Instruments, Sarasota, FL), and dental cement (Pentron Clinical, Orange, CA) was cured with a dental curing light to make a head-cap. Immediately after, EIS was measured, and tonic the 5-HT response was measured using SWV over a five-minute period. After surgery, animals were placed on an electric heating blanket to wake up and received an i.p. injection of 5 mg/kg ketofen (100 mg/mL, Zoetis Inc, Kalamazoo, MI, USA) daily for three days post-surgery. SWV detection was repeated every day for one week. SWV experiments were acquired using a potentiostat/galvanostat (Autolab PGSTAT128N, Metrohm, USA) connected in a three-electrode configuration: working electrode, bone screw counter electrode, and Ag/AgCl wire reference electrode. 5-HT peaks were isolated from the non-faradaic background current for each SWV scan by subtracting a modelled polynomial baseline using a previously described methodology (Taylor et al. 2019). The *in vivo* 5-HT concentration was determined for all *in vivo* experiments by converting the SWV peak current to 5-HT concentration using the pre-calibration electrode sensitivity, as previously reported (Castagnola et al. 2022; Taylor et al. 2019).

For the biocompatibility/histology study, 15 mice (3 groups of 5 mice at 3 end time points, i.e., one, two and four weeks) were bilaterally implanted with dummy flexible MEAs and dummy stiff silicon probes (nonfunctional 3-mm A-style probes, Neuronexus). Two electrode holes were drilled bilaterally over CA2 region of the hippocampus (AP: -2.91, ML: +/-3.35, DV: -2.5 to -3.0) (Paxinos and Franklin 2019), and the probes were inserted and sealed via Kwik-Cast Sealant (World Precision Instruments, Sarasota, FL, USA). Dental cement (Pentron Clinical, Orange, CA, USA) was used to anchor the probes in place. Incision sites were treated with a topical antibiotic, and animals received an intraperitoneal injection of 5 mg/kg ketofen (100 mg/mL, Zoetis Inc, Kalamazoo, MI, USA). Mice were placed back in their respective cages on top of a warming pad while they regained consciousness. Ketofen injections (5 mg/kg) continued daily for three days after surgery for pain management.

At one, two, or four weeks, animals were given an overdose of ketamine/xylazine (80−100 mg/kg and 5−10 mg/kg, respectively), which was confirmed by absent toe/tail pinch reflex. Mice were then intracardially perfused with 100 mL phosphate-buffered saline (PBS) followed by 100 mL 4% paraformaldehyde (PFA) in PBS. The bottom of the skull was removed to expose the brain and the skull was post-fixed in 4% PFA at 4 deg C for 4−6 h. Brains were sucrose protected, frozen, and cryosectioned at 25 μm thick slices.

### 2.6 Immunohistochemical staining

Immunohistochemical staining was performed on tissue slices containing flexible and stiff implant injuries; this method is consistent with previously published works from our group (Golabchi et al. 2018; Krahe et al. 2022). Frozen tissue sections were rehydrated in citrate buffer and then blocked with 10% goat serum and treated with 0.1% Triton-x for 45 min. Staining of the brains was performed in groups consisting of GFAP (DAKO rabbit 1:500) and tomato lectin (Vector Labs mouse 1:250). These stains allowed us to visualize reactive astrocyte migration to the implant (GFAP) and the presence of microglia as well as the intactness of the vasculature (tomato lectin). The intensity of GFAP and tomato lectin in the control (stiff probe) and experimental (flexible probe) groups, as well as the injury hole size left by the implants, were compared.

### 2.7 Image analysis

Images of slices were obtained using confocal fluorescent microscopy (Olympus Fluoview 1000) to evaluate the differences in tissue composition surrounding the implant sites; this method is consistent with previously published works from our group (Golabchi et al. 2018; Krahe et al. 2022). Images were analyzed in a custom MATLAB script. Bins of 10 µm were created concentrically around the probe implant and the intensity of the stain or number of cells labeled per bin was measured. Intensities and counts were scaled to control regions at the corners of the images. Background intensity was calculated from the corners of the image (20% of the total image area) by removing pixels greater than 1 standard deviation (STD) above the mean and calculating the mean and variance of the remaining pixels. All pixels greater than 1STD above the mean background intensity were used for analysis. The injury hole size was compared between groups via ImageJ, similar to a previously published method (Urdaneta et al. 2022). Manual selections were carefully drawn around the border of the implant holes and then the area within the selection was measured automatically via the Measure function. To normalize the size difference between stiff and flexible electrodes, a ratio between the selected hole area and the respective probe area (stiff: 1500 μm2, flexible: 2160 μm2). For week one, n = 15 control (stiff) images were compared to n = 18 experimental (flexible) images for both GFAP, tomato lectin, and injury hole size. For week two, n = 17 control images were compared to n = 20 experimental images. For week four, n = 15 control images were compared to n = 16 experimental images.

## 3. Results and Discussion

### 3.1 GC MEAs fabrication

GC MEAs were successfully fabricated using the high-resolution mask-less direct-writing photolithography process on SU-8 substrate that we previously reported (Castagnola et al. 2022), and schematically described in **Supplementary Figure 1**. This process enables the transfer of pre-patterned and pyrolyzed GC microstructures into a flexible substrate, enabling the integration of electrochemical detection capabilities, as previously demonstrated (Castagnola et al. 2022; Castagnola et al. 2021b; Castagnola et al. 2018). Combining the superior electrochemical stability of the carbon electrodes with the excellent biocompatibility of the thin-film flexible device (Fu et al. 2016; Luan et al. 2017; Xie et al. 2012), the GC MEA presents several advantages to achieve stable electrochemical sensing and seamless tissue integration.

Figure 1A-E shows front and side views of the fabricated MEA on SU-8 substrate with metal interconnections and GC microelectrodes. The device is composed of a singular shank (120 µm wide and ∼15 µm thick) with 6 circular GC electrodes 40 µm in diameter. An anchor hole at the tip has been patterned to facilitate the insertion of a 50 µm tungsten shuttle that enables the handling and penetration of the probes into the brain. The total length of the shank is 5.5 mm to target the CA2 region of the hippocampus of the mouse brain. GC MEAs are connected to the PCB using a zero-insertion force (ZIF) connector to interface with the potentiostat (Figure 1B). The flexibility of the MEA shank is demonstrated in Figure 1C, and the insertion of the tungsten shuttle guide is shown in Figure 1D. To analyze the chemical structures of the pyrolyzed carbon from SU-8, we collected the Raman spectra (Figure 1F). The Raman spectra shows the two primary peaks typical for carbon materials. The D band at 1323 cm^-1^ is due to breathing modes of six-atom rings, indicating defects from the boundaries like edge-plane or doping. The G band at 1604 cm^-1^ is due to E2g phonon mode, indicating sp2 graphitic carbon structures (Ferrari and Basko 2013; McCreery 2008). The Raman spectra also show some secondary peaks: the 2D peak at 2642 cm^-1^, which is the overtone of D peak, and the D+D’ peak at 2916 cm^-1^ (Ferrari and Basko 2013). The D/G ratio can be measured by calculating the peak height to evaluate the defect level of the carbon materials. The D/G ratio value is 1.18 for our pyrolyzed carbon from SU-8 100, which is comparable with the previously reported glassy carbon electrodes (Castagnola et al. 2021b). This result indicates that the pyrolyzed carbon from SU-8 are glassy carbon rich in defect sites that can be enhanced with functional groups to improve electrochemical properties (Cao et al. 2019).

### 3.2 PEDOT /CNT coated GC MEAs for SWV sensing of 5-HT tonic concentrations

Despite the outstanding electrochemical stability, GC electrode surface is smooth with limited effective surface area as shown in the optical images (Figure 2Ai and ii). Low surface area can lead to high electrode impedance and low sensitivity for SWV detection. High surface area PEDOT/CNT coating has shown significant effect in reducing impedance for electrophysiological recording (Alba et al. 2015; Kozai et al. 2016), electrical stimulation (Luo et al. 2011; Zheng et al. 2022), and increasing sensitivity for direct tonic DA measurements using SWV (Castagnola et al. 2022; Taylor et al. 2019). In this work we focus on optimizing PEDOT/CNT on GC and the SWV waveform for sensitive and selective detection of tonic 5-HT *in vivo*.

**Figure 2.**
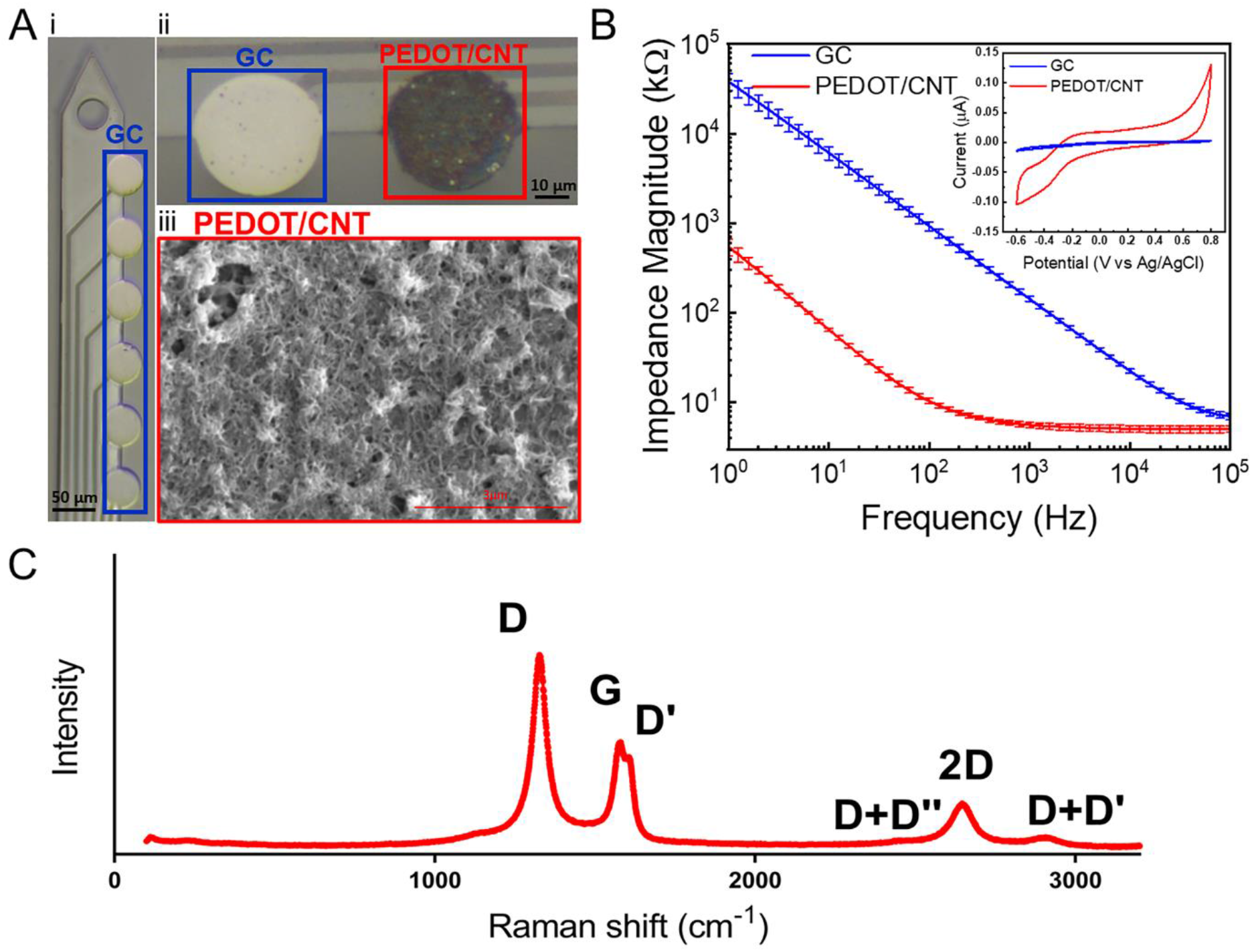
PEDOT/CNT coated GC-MEA. **A)** Optical picture of a GC MEA with GC microelectrodes (i). Magnification of 2 microelectrodes, 1 PEDOT/CNT coated GC in blue and 1 uncoated GC in red (ii). SEM image of the PEDOT/CNT high surface area structure (iii). **B)** Electrochemical impedance spectroscopy (EIS) plots of GC microelectrodes before and after PEDOT/CNT coating (n=6). In inset: representative cyclic voltammogram of PEDOT/CNT coated versus uncoated microelectrode. **C)** Raman spectra of PEDOT/CNT coatings.

The 40 µm diameter GC microelectrodes were successfully coated with PEDOT/CNT (Figure 2A ii, iii). A representative scanning electron microscopy (SEM) image of the PEDOT/CNT coating is reported in Figure 2Aiii. The high surface area structure (Figure 2A iii) of the PEDOT/CNT coatings drastically decreased the impedance over the entire measured frequency range from 143.68± 3.34kΩ to 5.65 ±0.40 kΩ at 1 kHz, (n=6), (Figure 2B). In the low frequency region (1-100 Hz) governed by the capacitive charge transfer, due to the increase of the surface area (Cogan 2008; Orazem and Tribollet 2008), the PEDOT/CNT coating reduced the impedance by two orders of magnitude (from 6.1 ± 1.1 MΩ to 65.3 ±2.8 kΩ at 10 Hz) (Figure 2B). The CVs of the PEDOT/CNT-coated GC microelectrodes confirmed an enhanced exchange of charges between the electrode and electrolyte due to the increased effective area of the nanostructured material (Figure 2B, inset). After the PEDOT/CNT coatings, the charge storage capacity (CSC), calculated as the time integral of an entire CV cycle divided by the geometric area, increased one order of magnitude, from 9.6 ± 1.6 mC/cm2 to 93 ± 10 mC/cm2. Qualitatively, the adhesion of PEDOT/CNT coating on GC is very stable, no delamination was observed after CV or SWV applications throughout this study. This is consistent with our previous results showing better adhesion between PEDOT and GC substrate than PEDOT on metal (Vomero et al. 2017) and very good adhesion of the PEDOT/CNT with the carbon substrates (Castagnola et al. 2017), even after prolonged electrical stimulation (Castagnola et al. 2017). Figure 2C shows the Raman spectra of PEDOT/CNT. The spectra show characteristic D and G peaks for carbon, and they also show the D’ peak due to double resonance (Ferrari and Basko 2013). The Raman spectra of PEDOT/CNT is the exact same as pure functionalized CNTs (**Supplementary Figure 2**), indicating PEDOT does not contribute to Raman responses.

### 3.3 *In vitro* SWV sensing of 5-HT tonic concentrations

The performance of PEDOT/CNT coated GC MEAs for the electrochemical detection of 5-HT was first determined via *in vitro* calibration experiments performed in aCSF, in the presence of several electrochemically active species that are likely also to be present *in vivo*: 3,4-dihydroxyphenylacetic acid (DOPAC, 10 µM), dopamine (DA, 1 µM), epinephrine (EP, 5 µM), 5-hydroxyindoleacetic acid (5-HIAA, 5 µM), ascorbic acid (AA, 1 mM) and uric acid (UA, 10 µM). The electrode sensitivity was determined based on the slope of the calibration curve (relating peak current to 5-HT concentrations) over a concentration range of 0.01-1μM. The SWV waveform for 5-HT determination was optimized by varying the parameters that can influence the voltammetric responses, i.e. frequency, step potential, pulse amplitudes, holding potential and holding time (Levent 2012). We found that the optimum waveform is a square wave with 50 mV pulse amplitude, 5 mV step height and 40 Hz frequency, scanned from 0.15 to 0.45 V. We chose to use 0.15 V as a lower bound and to hold the potential at 0.15 V in between scan repetitions to avoid electrochemical fouling due to reaction byproducts. In particular, a quinoneimine byproduct has been previously observed to be produced in 5-HT rich regions at around 0.05 V and results in significant electrode fouling shortly after 5-HT exposure (Castagnola et al. 2020b). Using these SWV conditions, the PEDOT/CNT microelectrodes show high sensitivity towards 5-HT (17.56 ± 0.01 nA/µM, n=5), 16-times higher than that of CFEs whose geometric area is 7 times larger (1.19 ± 0.01 nA/µM, n=5) (Figure 3 A-B). Using the 5-HT optimized waveform we observed excellent selectivity of the sensor towards 5-HT, as shown in Figure 3C, because the oxidation peaks of EP, DA, DOPAC, UA and AA are all outside of the 0.15–0.45 V potential window. By extending the potential window to -0.1V, we can see the oxidation peaks at 0.11V for EP, DA, DOPAC, 0.14V for AA, and 0.22 V for UA at relatively high concentrations (reported in Figure 3C). Only 5-HIAA has an oxidation peak overlapping with 5-HT. We tested the sensitivity of the PEDOT/CNT coated MEA toward 5-HIAA alone, and toward 5-HT in presence of 1µM 5-HIAA (Figure SI 3 Supplementary Information). For 5-HT the average sensitivity is 19.71 nA/µM and a linear response range is 100 nM - 1 μM. On the other hand, 5-HIAA peak cannot be reliably detected at concentrations below 1 µM. Therefore, despite the oxidation peak overlap, our sensor is more sensitive and selective for 5-HT over 5-HIAA at the physiologically relevant concentration range (Gomez-Merino et al. 2001; Saylor et al. 2019; Virkkunen et al. 1995).

**Figure 3:**
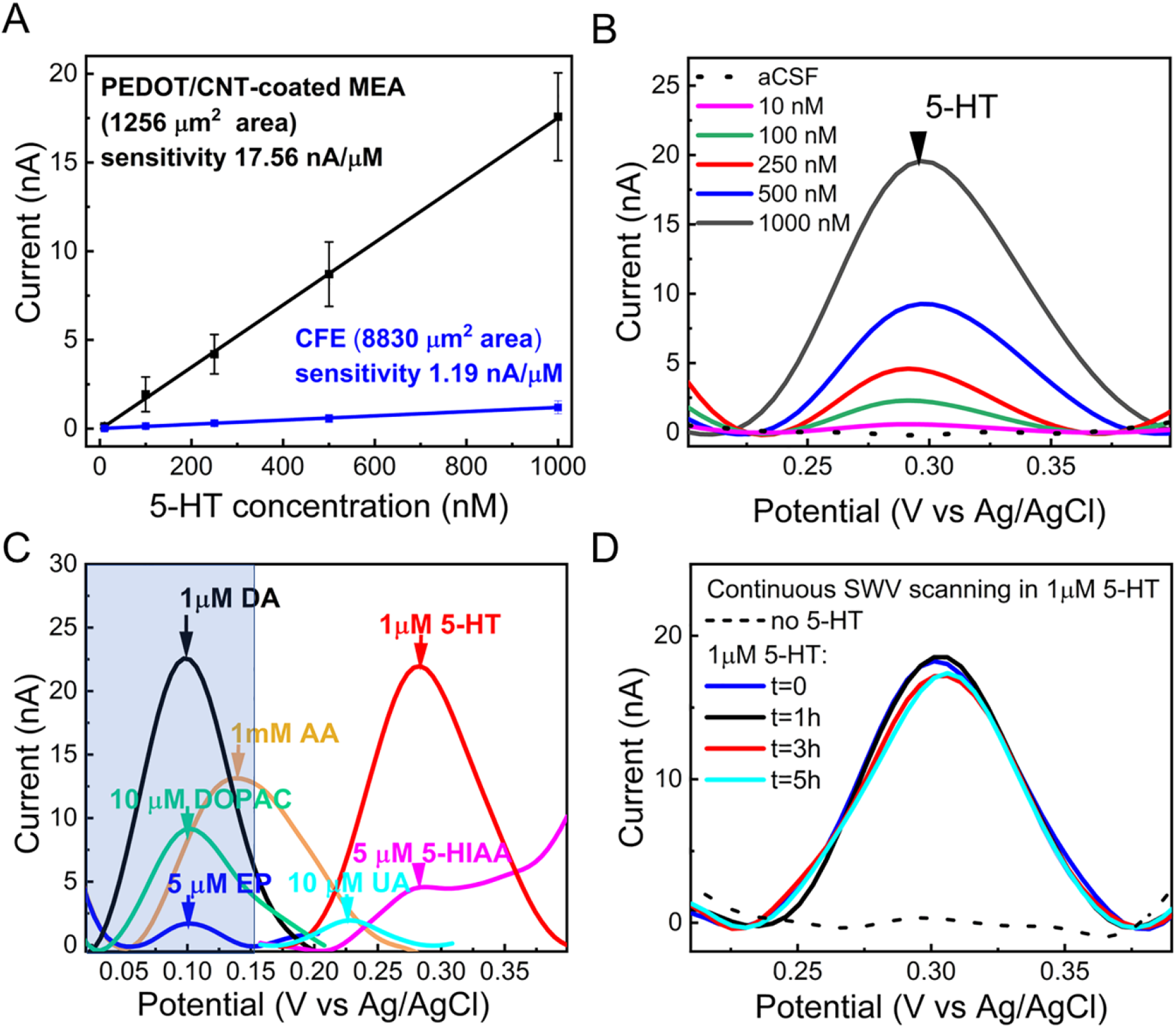
**A**) *In vitro* SWV 5-HT calibration plot *(peak current vs. 5-HT concentration, n=5)*, conducted at PEDOT/CNT coated GC MEAs (black) versus SWV 5-HT calibration plot conducted at CFEs with 7 times bigger geometric area, in aCSF in the presence of DOPAC, (10 µM), dopamine (1 µM), epinephrine (5µM), 5-hydroxyindoleacetic acid (5 µM), ascorbic acid (1mM) and uric acid (10 µM). The average sensitivity is linear in the range 10 nM - 1 μM and it is 16 times higher than the one obtained by CFEs with 7 times bigger geometric area. **B)** The corresponding SWV scans reveal clear 5-HT peaks at 0.29 V. **C)** Selectivity test showing a distinct 5-HT peak separated from the tested electrochemical interferants. **D)** Fouling test in the presence of 1 μM 5-HT (continuous SWV scanning): representative SWV in response to 1 µM of 5-HT at the time 0 and after 3, 4 and 5 h of continuous scanning.

The detection of 5-HT using electrochemical methods is challenging since the by-products formed during the oxidation reaction of 5-HT, such as reactive carbocation intermediate and dimers, are highly reactive and adsorb irreversibly on the electrode surface resulting in electrode fouling (Jackson et al. 1995; Patel et al. 2013; Wrona and Dryhurst 1990). Thus, to understand the fouling resistance of the PEDOT/CNT when exposed to 5-HT, we performed a fouling test corresponding to 5 hours of continuous SWV scanning in the presence of 1 μM 5-HT in aCSF. We observed good fouling resistance, with minimal variations in the peak current amplitude recorded over the 5h (Figure 3D). It is worth noting that the 5-HT concentration used in this fouling test is much higher than the physiological concentrations (Gomez-Merino et al. 2001; Saylor et al. 2019; Virkkunen et al. 1995).

### 3.4 *In viv*o SWV sensing of 5-HT tonic concentrations

The *in vivo* performance of PEDOT/CNT-coated GC MEAs for the detection of tonic 5-HT level was first determined through acute experiments conducted in the hippocampus of isoflurane-anesthetized mice. The flexible MEAs were implanted into the CA2 region of the hippocampus using a 50 µm tungsten wire inserted into the anchor hole at the shank tip (Figure 1A and D) and removed immediately after implantation, as previously reported (Castagnola et al. 2022). Immediately after, the tonic 5-HT response was detected using the optimized SWV waveform described above over a 20 min period. The *in vivo* SWV reveals clear 5-HT peaks at 0.29 V. Following 20 min of baseline data collection, the 5-HT peak was validated using stereotaxic direct infusion of 1 μL boluses of 5 µM 5-HT and 10 µM 5-HT solutions, to confirm the chemical specificity of our measurements (Figure 4A). SWV peak currents at 0.29 V vs Ag/AgCl (average and STD, n=6 mice) was converted to concentrations using the corresponding pre-calibration curves which resulted in a detected basal level of 5-HT of 77 ± 12 nM in the CA2 hippocampal region, in agreement with values previously detected in in the CA2 region of the mouse hippocampus (64.9 ± 2.3 nM) using FSCAV (Abdalla et al. 2017). Pre-calibration as opposed to post-calibration was used, similar to our previous reports (Castagnola et al. 2020a; Castagnola et al. 2022) due to the *in vivo* current response stability observed at PEDOT/CNT-functionalized electrodes. A drift-free current response is observed immediately after the MEA implantation without requiring electrochemical *in vivo* stabilization. On the other hand, we previously observed a high degree of coverage of the PEDOT/CNT-coated electrodes by biological matter (blood) upon explantation, which can induce a drastic decrease (>70%) in DA sensitivity *in vitro* measured in post-calibration experiments, potentially overestimating the *in vivo* concentrations.

**Figure 4:**
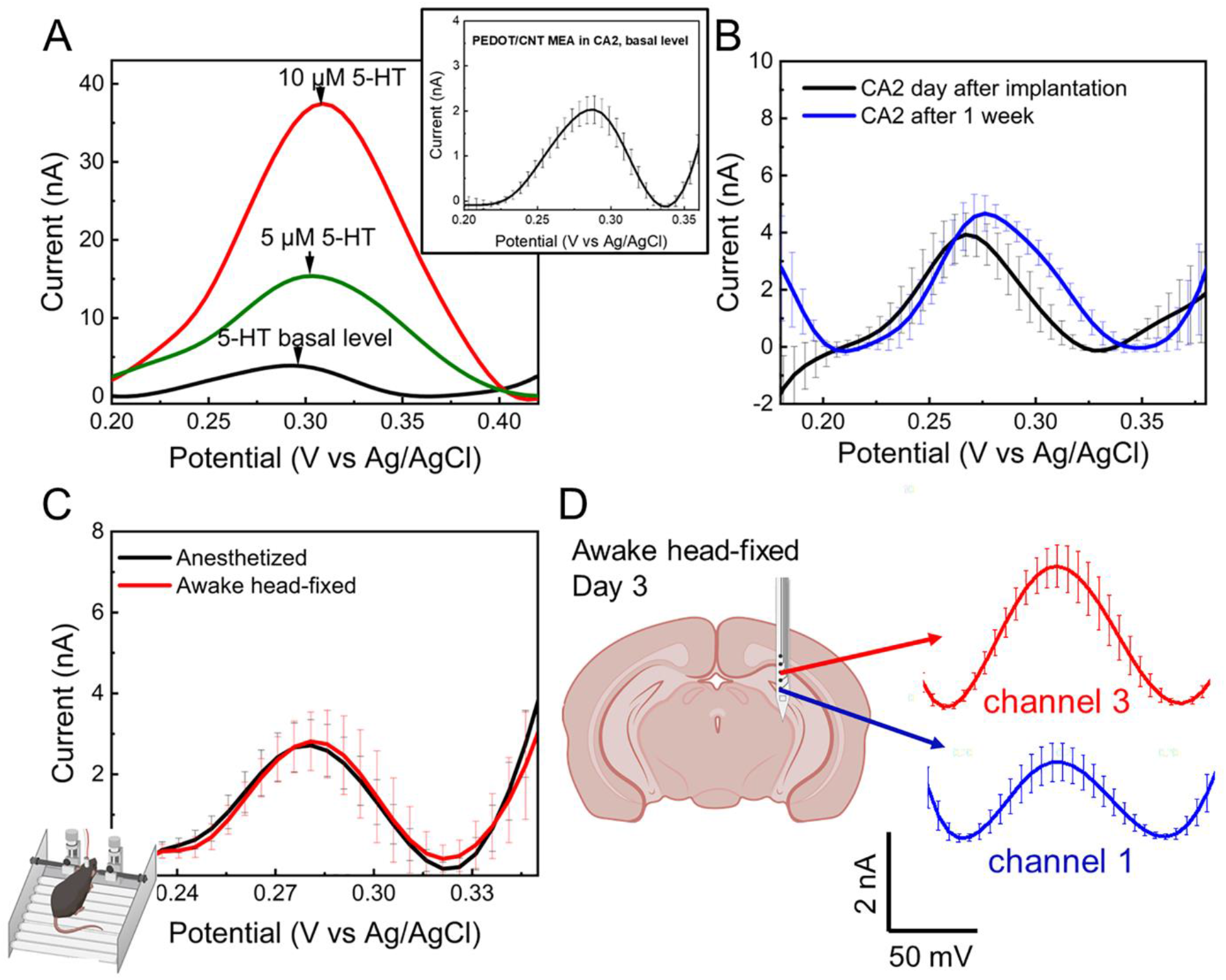
*In vivo* sensing in CA2 of mouse hippocampus. **A)** *In vivo* SWV conducted at PEDOT/CNT coated GC in the CA2 region of hippocampus (black) reveal clear 5-HT peaks at 0.29 V. Validation of the 5-HT peak using stereotaxic direct infusion of 1 μL boluses of 5 µM 5-HT (green) and 10 µM 5-HT (red) solutions. Higher concentrations resulted in larger 5-HT peak currents, with a slight shift in potential, possibly due to pH. In inset: magnification of the 5-HT basal peak (average and standard deviation, n=6 mice) **B)** *In vivo* 5-HT SWV detections at PEDOT/CNT coated MEAs one day (black) and one week (blue) post-implant show that the sensor can detect tonic 5-HT for at least seven days. **C)** *In vivo* SWV conducted at PEDOT/CNT coated MEA in the CA2 region of hippocampus of anesthetized versus awake head-fixed mouse show similar 5-HT peaks, demonstrating the feasibility of our SWV technique for awake mice. **D)** Representative *in vivo* SWV 5-HT detections at different channels of the same PEDOT/CNT coated MEA after three days of implantation, showing the feasibility of chronic multichannel recording in awake head-fixed mice.

To test the *in vivo* stability of the 5-HT tonic detection, as well as the capability of our MEAs to perform in head-fixed awake mice, we chronically implanted the PEDOT/CNT coated GC MEAs in the CA2 hippocampal region of 3 mice. Figure 4B shows the *in vivo* 5-HT SWV detection at PEDOT/CNT coated MEAs at one day (black) and one week (blue) post-implant in anesthetized mice, demonstrating that the PEDOT/CNT-coated MEA can stably detect tonic 5-HT for at least seven days. Though the peak amplitude is consistent along the first week of implantation, we observed a 0.03 V anodic shift in the oxidation peak during chronic experiments, which is likely caused by the de-chlorination of the chronically implanted Ag/AgCl reference electrodes, resulting in errors in the potential reading at the working electrode (Robbins et al. 2022). Subcutaneous Ag/AgCl reference freshly inserted at each day of measurement will solve this problem (Robbins et al. 2022).

Despite the different efforts reported in waveform optimization to reduce the fouling at CFEs (Jackson et al. 1995; Shin et al. 2021; Yoonbae Oh 2016), to the best of our knowledge, this is the longest *in vivo* 5-HT detection ever reported. This is a significant advance, as electrochemical fouling is a well-reported obstacle for long term 5-HT detection (Jackson et al. 1995). The oxidation reaction of 5-HT involves a two-electron, two-proton transfer process (Jackson et al. 1995; Ou et al. 2019; Patel et al. 2013), during which byproducts such as reactive carbocation intermediates and dimers are formed (Jackson et al. 1995; Patel et al. 2013; Verbiese-Genard et al. 1984). These byproducts can adsorb irreversibly on the electrode surface resulting in electrochemical fouling (Jackson et al. 1995; Wrona and Dryhurst 1990), reducing or destroying detection sensitivity. Using our optimized waveform that starts scanning at 0.15V up to 0.45 V and holds the potential at 0.15 V in between scans, we minimize the electrochemical fouling due to secondary reaction byproducts; in particular, the quinoneimine byproduct that has been previously been observed to be produced at around 0.05 V(Castagnola et al. 2020b). Using these SWV conditions, the PEDOT/CNT-coated GC microelectrodes achieve excellent 5-HT selectivity, avoiding the potential range at which EP, DA, DOPAC, UA and AA oxidize. The fouling resistance to 5-HT reaction byproduct may also be improved by the incorporation of the negatively charged acid functionalized CNTs into the PEDOT/CNT coating, similar to what was previously observed for CNT-based microelectrodes (Mendoza et al. 2020; Zestos et al. 2014) and high surface area carbon materials (Mendoza et al. 2020; Swamy and Venton 2007; Weese et al. 2019; Yang et al. 2017; Zestos et al. 2014). Their antifouling properties have been attributed to the presence of defect sites in high density at their edge and electrocatalytic-active functional groups that can increase the surface hydrophilicity and reduce adsorption of molecules (Castagnola et al. 2021a; Garg et al. 2017; Puthongkham and Venton 2020). These findings are also supported by the Raman spectra (Figure 2C). The Raman spectra of the functionalized CNTs show high D/G peak ratio, indicating a large portion of edge-plane sites. We then compared the 5-HT sensing performance of our sensor in anesthetized versus awake head-fixed mice and found similar 5-HT peaks in SWV measurements, demonstrating the feasibility of our SWV technique for awake mice. Figure 4C presents a representative example for *in vivo* SWV 5-HT detection, collected at day three post implantation from an awake head-fix mouse at two different channels of the same PEDOT/CNT coated GC MEA.

Regardless of tonic or phasic detection, there has been no report of multisite measurements of 5-HT from the brain. The capability of stable multichannel measurements opens the possibility to map the 5-HT dynamics in different brain regions or different loci of the same region. Additionally, because GC microelectrodes have already demonstrated to provide high sensitivity FSCV detection of 5-HT (Castagnola et al. 2021b); the GC MEAs have the unique potential to combine the detection of both phasic and tonic 5-HT dynamics over multiple timescales (milliseconds to minutes) at multiple brain locations, both in anesthetize and awake mice.

### 3.5 *In vivo* biocompatibility: flexible GC-MEAs versus stiff probes

Several studies, mainly conducted in the cortical region of rodents, have shown that reduced probe shank cross sectional area and mechanical mismatch between probe and brain tissue contribute to successful chronic tissue integration and improved recording longevity (He et al. 2020; Luan et al. 2017; Vomero et al. 2022; Xie et al. 2012; Zhao et al. 2019). However, to detect 5-HT the flexible MEAs need to be chronically implanted in the hippocampus, a region especially vulnerable to damage caused by metabolic dysregulation or oxidative stress (Jackson and Foster 2009; Kirino et al. 1985; McEwen 1994). Here, we provide immunohistochemistry analysis to study host tissue response to our flexible MEAs implanted in the CA2 region of the hippocampus over a four-week period, in comparison with stiff commercially available neural probes, with similar thickness (15µm) and 20% smaller width (120 vs. 80µm). Intermediate end points (7 and 14 days) were strategically chosen to provide histological and explant snapshots to characterize the time course of probe-induced immune response.

Neural tissue slices from animals implanted bilaterally with stiff dummy probes and our experimental flexible probes were compared based on reactive astrocyte presence (GFAP), microglia encapsulation and vascular presence (tomato lectin), and the injury site size. We used GFAP to determine the astrocytic activation around the electrode, while tomato lectin was used to stain microglia and tissue vasculature. The packing of these astrocytes and microglia around the electrodes has shown to directly affect signal recording, sensing ability, and stimulation quality (Kozai et al. 2015; Xu et al. 2019). One week after implantation (Figure 5), there is a significant difference (p < 0.001, Wilcoxon signed rank test) in the tomato lectin expression surrounding the stiff versus flexible electrodes, especially within the first 50 μm, where microglia are expected to accumulate (Figure 5H) (Bjornsson et al. 2006; Kozai et al. 2015). This translates to an increase in microglia and vascular remodeling in the brain tissue near the stiff implants compared to the flexible implants, despite a 40% wider shank of the flexible probes. Observing the lack of intact blood vessels in the image window of week one versus weeks two and four suggests that the injured tissue has not had sufficient time at one week to clear out broken vessels while regenerating lost ones, in addition to larger amounts of microglia moving towards the probe in the first week. This is supported by Supplemental Figure 4, which represents the untouched tissue within the same animals, and clearly displays intact blood vessels similar to those seen near the week two and four tomato lectin images. A higher mean GFAP intensity was found in the first 50 um bin from the implant, but the differences in GFAP intensity between the two groups were not statistically significant (Figure 5G).

**Figure 5.**
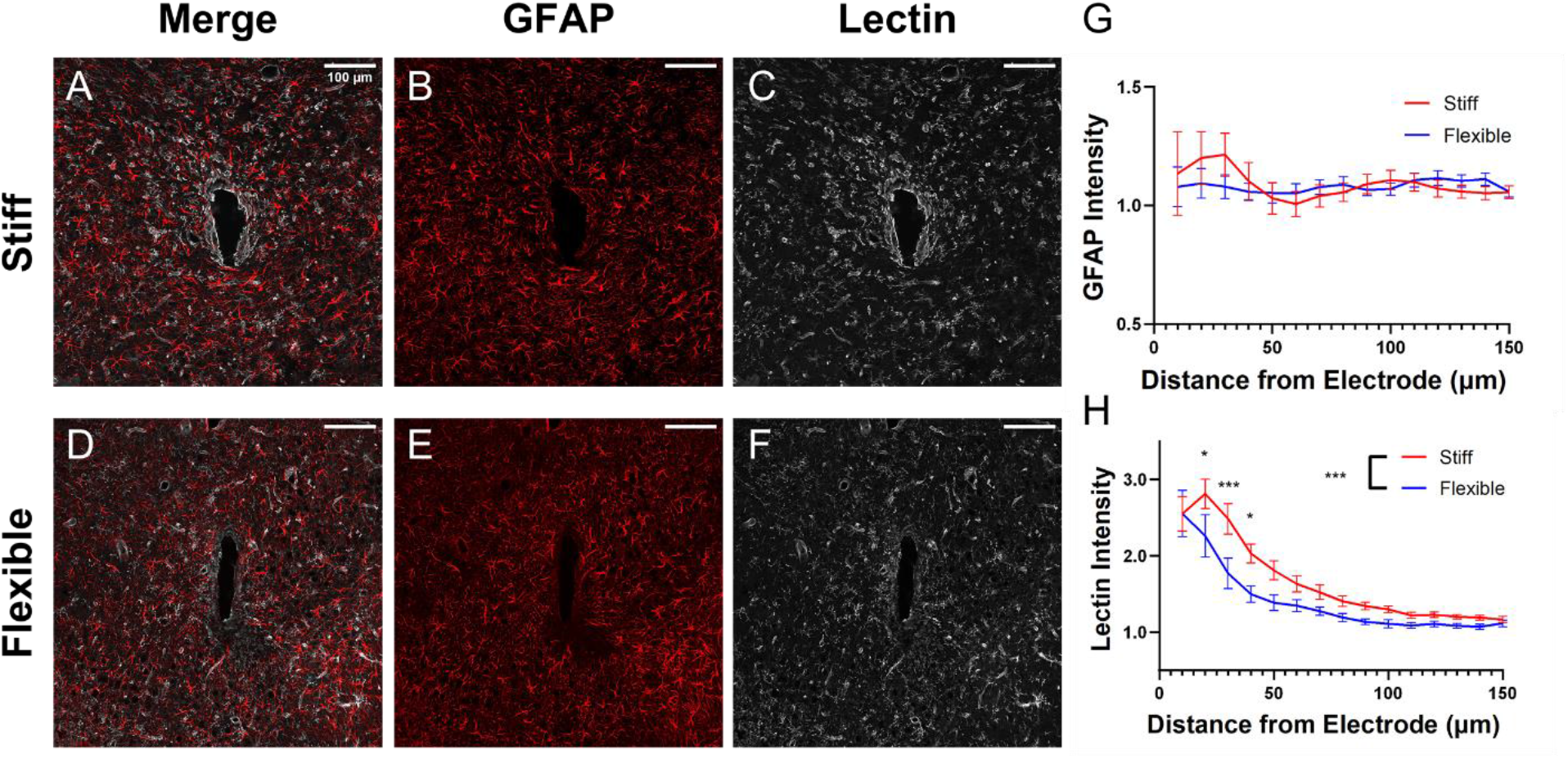
Microglia accumulation and vascular remodeling after one week caused by stiff and flexible probes. Merged GFAP and tomato lectin images of **A)** stiff and **D)** flexible probe scars. GFAP channel only for **B)** stiff and **E)** flexible probe scars. Tomato lectin channel only for **C)** stiff and **F)** flexible probe scars. **G)** GFAP intensity over several distances from the probe center. **H)** Tomato lectin intensity over several distances from the probe center (Wilcoxon signed rank test, * = p < 0.05, *** = p < 0.001).

For brain tissue analyzed two weeks after implantation (Figure 6), significant differences were found between stiff and flexible probe groups in both GFAP and tomato lectin expression. GFAP intensity (indicating reactive astrocyte response) surrounding the stiff electrodes was significantly higher overall than the intensity surrounding the flexible implants (Figure 6 G) (p < 0.01, Wilcoxon signed-rank test). Additionally, tomato lectin intensity and thus microglia/vascular remodeling presence remained significantly increased overall in the stiff group (p < 0.0001), again with the most significant differences occurring within the first 50 μm (Figure 6H) (p < 0.0001 at 10 μm, p < 0.01 at 20 μm).

**Figure 6.**
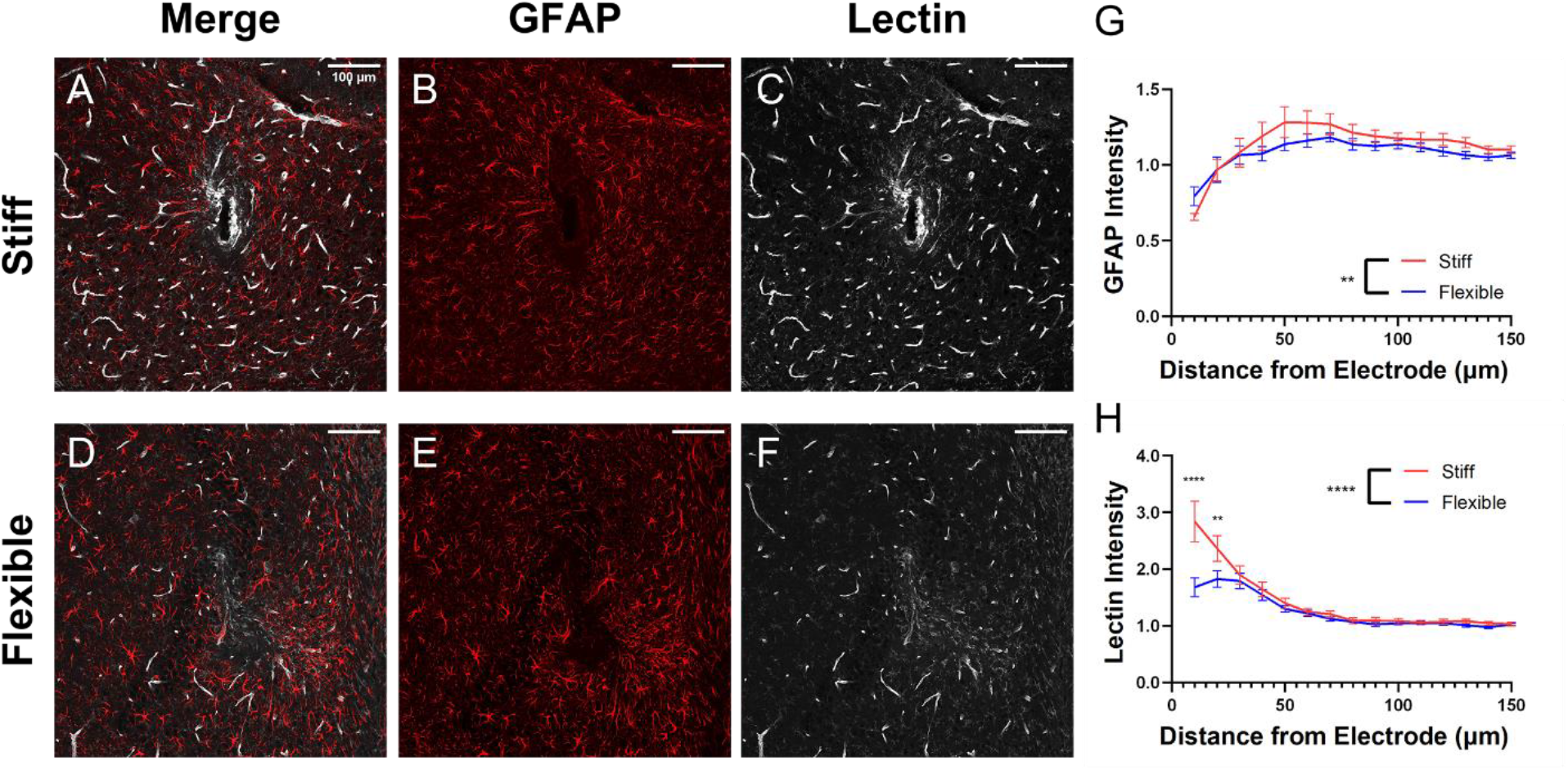
Blood vessel injury, and astrocyte and microglia accumulation in two weeks caused by stiff and flexible probes. Merged GFAP and tomato lectin images of **A)** stiff and d) flexible probe scars. GFAP channel only for **B)** stiff and **E)** flexible probe scars. Tomato lectin channel only for **C)** stiff and **F)** flexible probe scars. **G)** GFAP intensity over several distances from the probe center. **H)** Tomato lectin intensity over several distances from the probe center (Wilcoxon signed rank test, ** = p < 0.01, **** = p < 0.0001).

At four weeks, significant differences between the tissue surrounding stiff and flexible electrodes were sustained (Figure 7). GFAP intensity was significantly greater in the flexible group (Figure 7G, p < 0.01, Wilcoxon signed rank test), with the average value remaining close to one (which indicates no increase compared to control regions far from the electrode). While tissue immediately near the stiff electrodes expressed significantly less GFAP, it should be noted that the average hole size created by the stiff implants was significantly higher than for flexible electrodes in week four, which may have heavily contributed to the low (less expression than far from the electrode) average GFAP expression close to the stiff implants (Figure 8F). The trend for tomato lectin expression is consistent with weeks one and two, with intensity significantly higher surrounding the stiff electrodes than our flexible electrodes (Figure 7H) (p < 0.0001).

**Figure 7.**
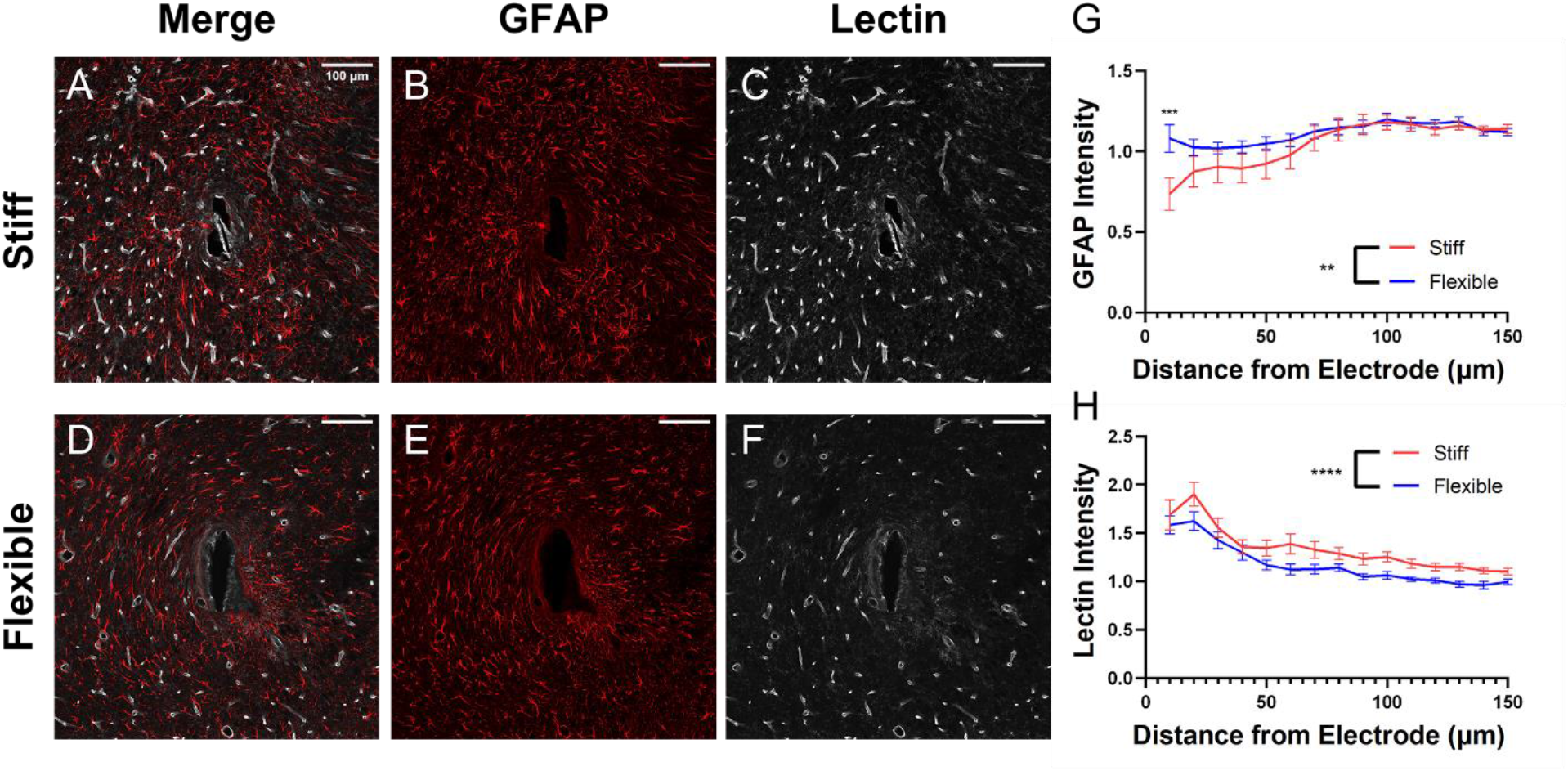
Microglia accumulation and vascular remodeling after four weeks caused by stiff and flexible probes. Merged GFAP and tomato lectin images of A) stiff and D) flexible probe scars. GFAP channel only for B) stiff and E) flexible probe scars. Tomato lectin channel only for C) stiff and F) flexible probe scars. G) GFAP intensity over several distances from the probe center. H) Tomato lectin intensity over several distances from the probe center (Wilcoxon signed rank test, ** = p < 0.01, *** = p < 0.001, **** = p < 0.0001).

**Figure 8.**
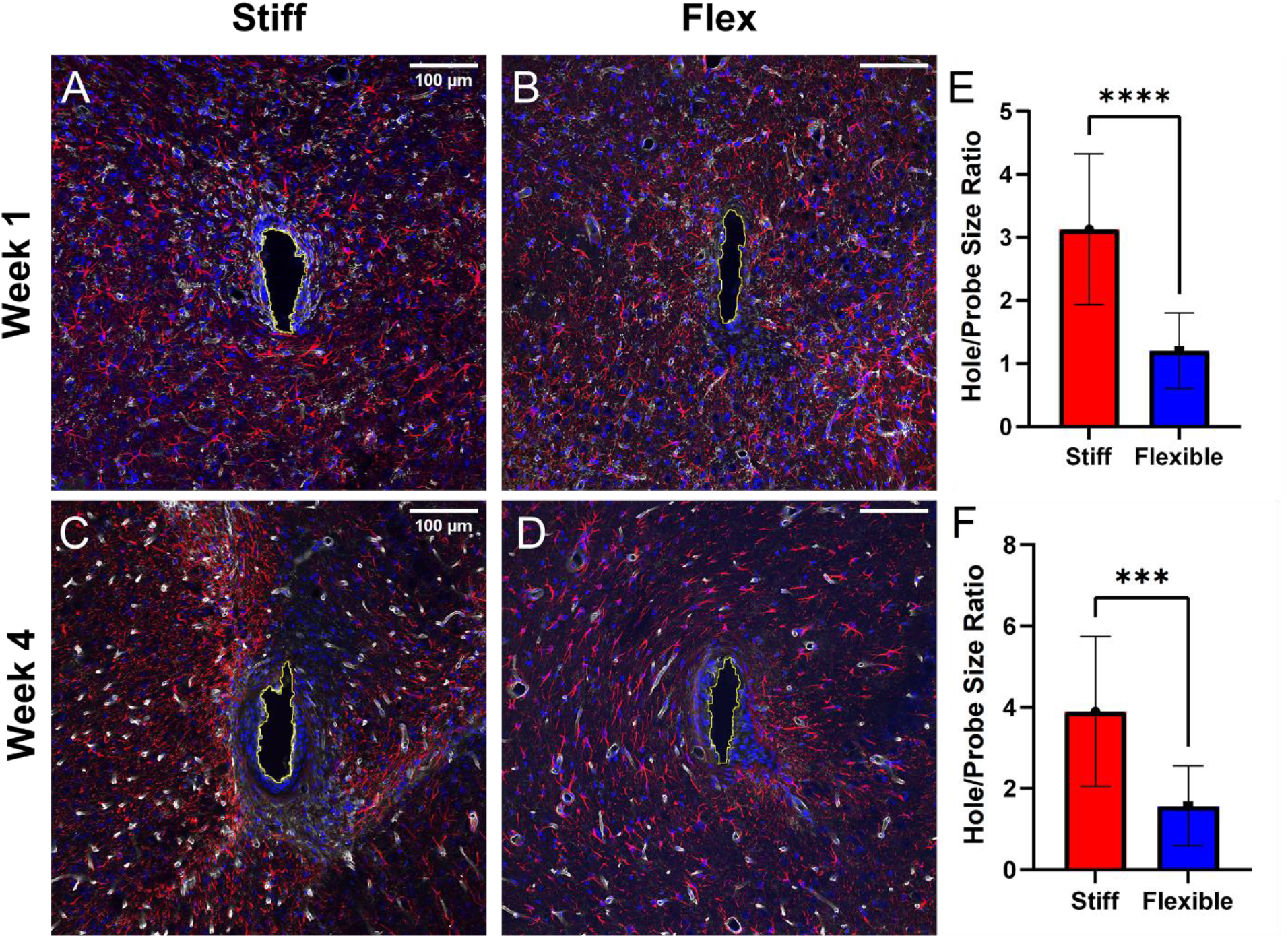
The explant hole size was decreased in acute and chronic setups by using a flexible probe. **A)** DAPI, GFAP, tomato lectin overlay for stiff implant after one week. A manually drawn explant hole is shown in yellow. **B)** Overlay for flexible implant after one week. **C)** Overlay for stiff implant after four weeks. **D)** Overlay for flexible implant after four weeks. **E)** Hole size comparison between stiff and flexible after one week. The hole size was calculated by dividing the manually obtained explant hole by the respective probe area for stiff or flexible. **F)** Hole size comparison between stiff and flexible after four weeks. (*** = p < 0.001, **** = p < 0.0001, Wilcoxon signed-rank test).

It was necessary to compare the size of the explant hole created by the stiff or flexible probe as an indication of tissue damage by the implant itself and the micromotion. In both weeks one and four, the stiff electrode created a significantly bigger hole than the flexible electrodes, despite the 40% smaller width. The hole size was normalized by the cross-section area of the device and represented as ratios. The stiff implants created a hole with a 2.60x larger average ratio in week one and a 2.48x larger ratio in week four (Figures 8 E and F). Overall, the flexible electrodes left a smaller footprint in the surrounding neural tissue, this is likely due to the fact that the flexible electrode can move with the neural tissue and mitigate tears due to micromotion. Reduced micromotion will also lead to reduced inflammatory response as observed in our GFAP and tomato lectin analysis.

In summary our histological analysis showed that flexible MEAs lead to better preservation of surrounding neural tissue and reduced astrocyte and microglia encapsulation. This finding is important because the recording, sensing, and stimulation quality of the electrodes is directly impacted by the astrocytic and glial scarring that accumulates around the electrodes (Kozai et al. 2015). Intimate contact with healthy neural tissue is tremendously important for sensing applications. A physical gap, barrier, or an area of damaged tissue between healthy serotonergic neurons and the detecting electrode will hinder diffusion of 5-HT to the electrode surface and distort the measurement. With minimum micromotion, more intimate integration and a reduced inflammatory response, the flexible electrodes are poised to provide stable sensing performance as observed here. This result points out that the use of flexible substrates plays a more significant role than the probe size, in reducing the FBR and improving chronic tissue integration. This aspect should be further investigated in future studies, using flexible GC MEAs with different cross-sectional areas.

GC microelectrodes already demonstrated excellent performance in 5-HT phasic sensing using FSCV. Here, we further demonstrate their capability to detect tonic 5-HT levels in anesthetized and awake mice. Overall, the results of this study suggest that our flexible GC MEAs represent a promising platform for investigating the role of phasic and tonic 5-HT dynamics in various brain functions and dysfunction as well as studying mechanisms of action of therapeutic treatments.

## 4. Conclusions

Here we developed the first implantable sensor tested *in vivo* for multi-channel 5-HT tonic detection in the hippocampus of mouse brain. Batch-fabricated GC-MEAs on flexible SU-8 substrates using the high-resolution mask-less direct-writing photolithography process produced highly reproducible MEAs with high quality GC microelectrodes. The optimized PEDOT/CNT coatings and SWV waveform resulted in a) exceptional sensitivity and selectivity toward 5-HT, and fouling resistance *in vitro*, and b) multichannel detection of tonic 5-HT concentrations *in vivo*, in the hippocampus (CA2) of mice in both anesthetized and awake head-fixed conditions. Notably, we demonstrate here that our flexible PEDOT/CNT-coated GC MEAs microelectrodes can achieve selective 5-HT tonic detection in the mouse hippocampus (CA2) for one week, the longest recording ever reported. Quantitative immunohistochemical analysis reveals reduced tissue damage and inflammation compared to stiff silicon probes with similar thickness and 40% smaller width, proving a healthy neural tissue interface that can facilitate long term sensing measurements Because GC microelectrodes have already demonstrated to provide high sensitivity FSCV detection of 5-HT (Castagnola et al. 2021b) and high quality electrophysiological recordings, our results highlight the potential of flexible GC MEAs as a platform for integrated multi-channel neurochemical and electrophysiological recording, a key tool to understand brain functions and neurochemistry. Future work will focus on integrating multiple modalities in one device, i.e. as neurochemical sensing over multiple timescales (milliseconds to minutes) and electrophysiological recording.

## Supporting Information

The Supporting Information is provided as a separate file.

## Author Contributions

EC and XTC conceptualized the project. EC fabricated the GC MEAs, performed the electrochemical characterization and sensing experiments *in vitro* and *in vivo*, performed *in vitro* fouling experiments and data analysis, and wrote the manuscript. EC performed surgeries for the Biocompatibility study. EMR perform part of the in vivo experiments and perfusions. EMR and DK and performed the biocompatibility study of flexible GC-MEAs versus stiff probes. DK wrote the biocompatibility study section of the manuscript. B.W., M.Y.P and QC performed the GC pyrolysis and fabricated part of the GC microelectrode arrays. Q.C. performed Raman characterization. B.W. performed SEM imaging. XTC supervised the fabrication, and device characterization, supervised the sensing experiments, and edited and revised the manuscript. All authors contributed to the article and approved the submitted version.

## Conflict of Interest Contributions

All authors have no conflict of interest to declare.

