## Supplementary Information for "Implantable flexible multielectrode arrays for multi-site sensing of serotonin tonic levels"

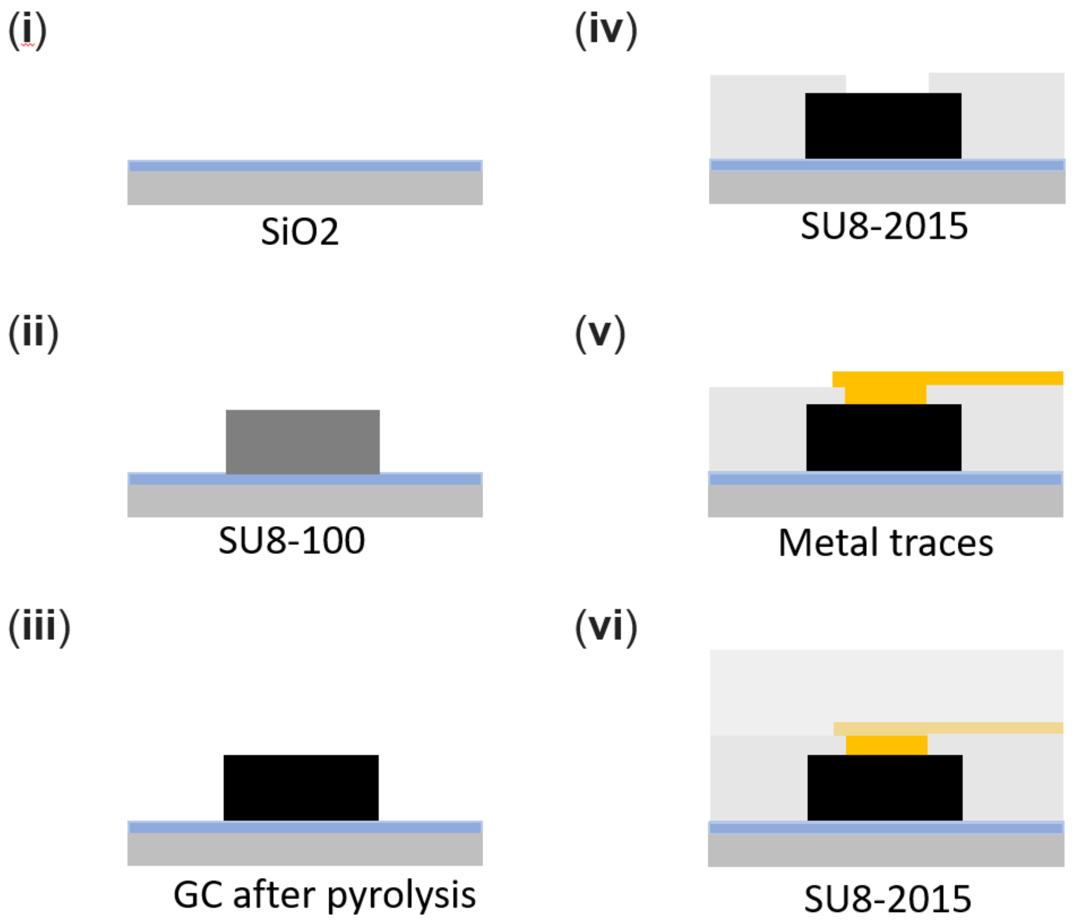


**SI Figure 1. Schematic for fabrication of the GC MEAs:** (i) (ii) SU-8 100 spin-coating and patterning of electrodes on $SiO_2$ wafer; (iii) pyrolysis; (iv) SU-8 insulation layer spin-coating on top of the GC electrodes and UV exposure to pattern the insulation layer and open a connection between the GC electrodes and the metal traces (next step) and the anchor hole; (v) Metal deposition and patterning using a lift-off procedure; (vi) SU-8 top insulation layer spin-coating and patterning of the probe outline and the anchor hole for the insertion of a 50 µm tungsten shuttle. Finally, the probes were released from the silicon substrate using buffered oxide etchant (1:7) in acid hood.

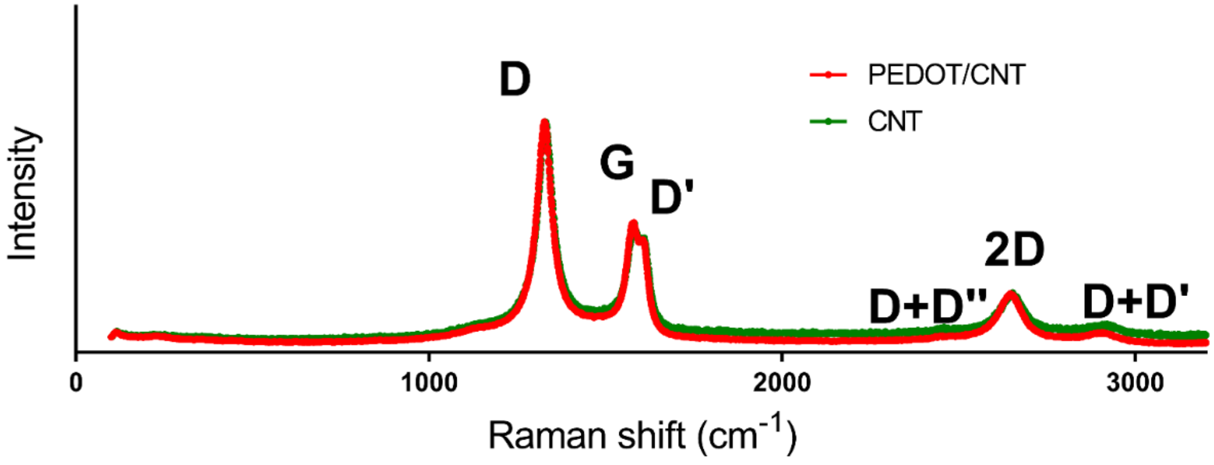


**SI Figure 2.** The Raman spectra comparison of PEDOT/CNT and CNTs.

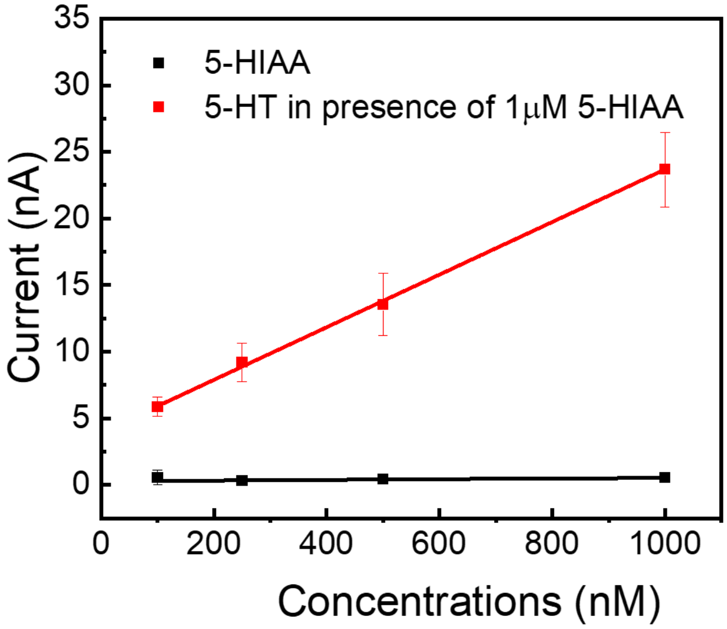


**SI Figure 3.** *In vitro* square wave voltammetry (SWV) calibration plot (peak current vs. concentration, n=5), conducted at PEDOT/CNT coated GC MEAs in aCSF toward 5-HIAA alone (black) and toward 5-HT (red) in presence of 1µM 5-HIAA. For 5-HT the average sensitivity is 19.71 nA/µM in the linear in the range 100 nM - 1 µM. On the other hand, 5-HIAA peak cannot be reliably detected at concentration at concentration <1 µM.

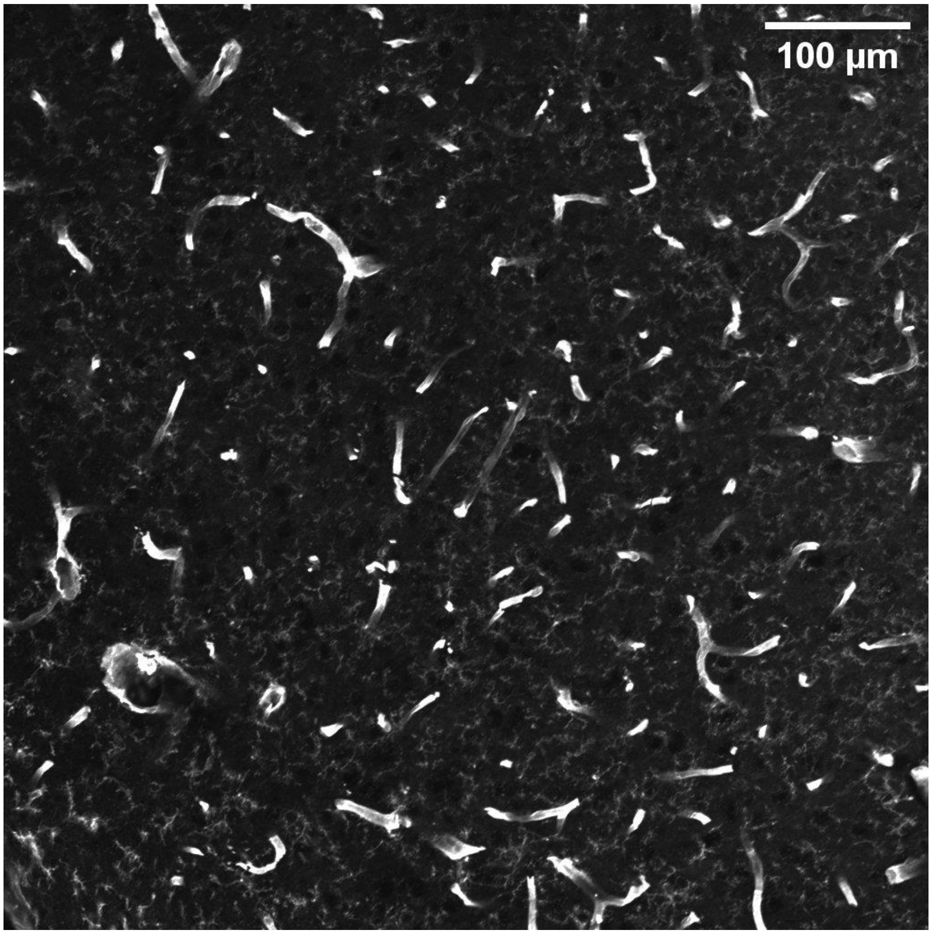


**SI Figure 4** Vasculature far from implant sites in week one animals remain intact.
